# *FOXS1* is a Master Regulator of Pathological Epithelial to Mesenchymal Transition in Human Epithelia

**DOI:** 10.1101/154369

**Authors:** Timothy A Blenkinsop, Tomasz Swigut, Nathan Boles, Rajini Srinivasan, Alvaro Rada-Iglesias, Qingjie Wang, Jeffrey H Stern, Joanna Wysocka, Sally Temple

## Abstract

Epithelial to mesenchymal transition (EMT) is a biological process involved in normal tissue morphogenesis and also in disease pathology. It causes dramatic alterations in cell morphology, migration, proliferation, and phenotype. We captured the global transcriptional and epigenetic programs elicited when polarized, cobblestone human retinal pigment epithelial (RPE) cells were stimulated to undergo EMT, a process associated with several retinal pathologies. The reorganization of chromatin landscapes occurred preferentially at distal enhancers, rather than promoter regions, accompanied by 3136 significantly changing genes. Of the 95 significantly changing transcription factors, FOXS1 was most upregulated. Loss and gain of function experiments demonstrated that FOXS1 is upstream of canonical EMT regulators, and can stimulate EMT in RPE and other epithelia. Inhibition of p38, stimulated by combined action of the EMT-inducing factors, TGFβ1 and TNFα, dampened *FOXS1* expression. An increase of *FOXS1* in several cancers indicates it has a role in several EMT-involved pathologies.

## Introduction

During EMT, epithelial cells change phenotype, lose cell polarity and typically become migratory and proliferative ^1^. Several pathological conditions involve EMT-related processes resulting in cell invasion and metaplasia. In the retina, the retinal pigment epithelium (RPE) forms a single, non-proliferative epithelial cell layer that supports neural retinal functions. RPE cells are polarized and are connected by tight junctions that form the blood-retina barrier. In pathological conditions, the RPE can undergo EMT and delaminate, migrating through the retina into the vitreous. There the cells proliferate and acquire mesenchymal characteristics, leading to epiretinal membrane formation, such as seen in proliferative vitreoretinopathy (PVR), the most damaging source of retinal detachment and vision loss in developed countries ^2^. While many factors have been associated with RPE changes in PVR, the critical factors and the changes they elicit in RPE that lead to its contribution in PVR are unknown.

We found that *TGFβ1* and *TNFα* pathways synergistically activate an EMT program in adult human RPE, producing changes similar to those observed in PVR. To characterize the molecular mechanism underlying this cellular transition, we mapped epigenomic and transcriptional changes and identified a set of transcription factors that are upregulated upon EMT induction in the adult RPE. Among those, we found that *FOXS1*, while not expressed in normal RPE, is highly induced in response to *TGFβ1* and *TNFα* and, notably, is present in patient PVR biopsy. We found that *FOXS1* is upstream of canonical regulators of the EMT program, *SNAIL* and *SLUG*, in RPE cells. Furthermore, consistent with a more general role in promoting EMT in various biological contexts, *FOXS1* also induced *SNAIL* expression in human mammary cells and activated *SLUG* and *TWIST* expression in hepatic epithelial cells, driving them towards EMT. Finally, increased copy number of the *FOXS1* gene locus is found in numerous metastatic cancer studies. These observations identify *FOXS1* as a master regulator of EMT that could play a driving role in several pathological contexts.

## Results

*TGFβ1* and *TNFα* have long been known to be present in the vitreous of patients with PVR ^3^ and both gene loci have SNP risk associations with PVR ^4,5^. Hence, we hypothesized that when applied to human RPE, their combination may model the PVR disease state.

RPE cells were isolated and expanded from adult human cadaver globes, then differentiated into a quiescent polarized cobblestone monolayer ^6^ (Fig. 1A). The resulting cobblestone RPE cells at passage 0 were treated with RPE with *TGFβ1* + *TNFα* (TNT) for 5 days, and gene expression tested by qPCR on RPE lines from three different donors, with measurements reported as mean ± SEM. TNT treatment led to the synergistic upregulation of EMT master transcription factors *SNAIL*, *SLUG* and *TWIST* (Fig. 1B). SNAIL expression increased 112.23 ± 27.91 (fold over control, set at one), *P* < 0.01 (*n=3*), after TNT treatment, whereas *TGFβ1* alone increased by 7.89 ± 2.21, *P* < 0.05 (*n=3*), and the *TNFα* change was not significant. SLUG expression increased 36.08 ± 6.31, *P* < 0.01 (*n=3*), in TNT treatment, while *TGFβ1* alone and *TNFα* alone had little effect. TWIST expression increased 11.26 ± 1.05, *P* < 0.01 (*n=3*), in TNT compared to control, while *TGFβ1* increased *TWIST* expression by 6.36 ± 1.91, *P* <0.05 (*n=3*), and *TNFα* had little effect.

**Figure 1.**
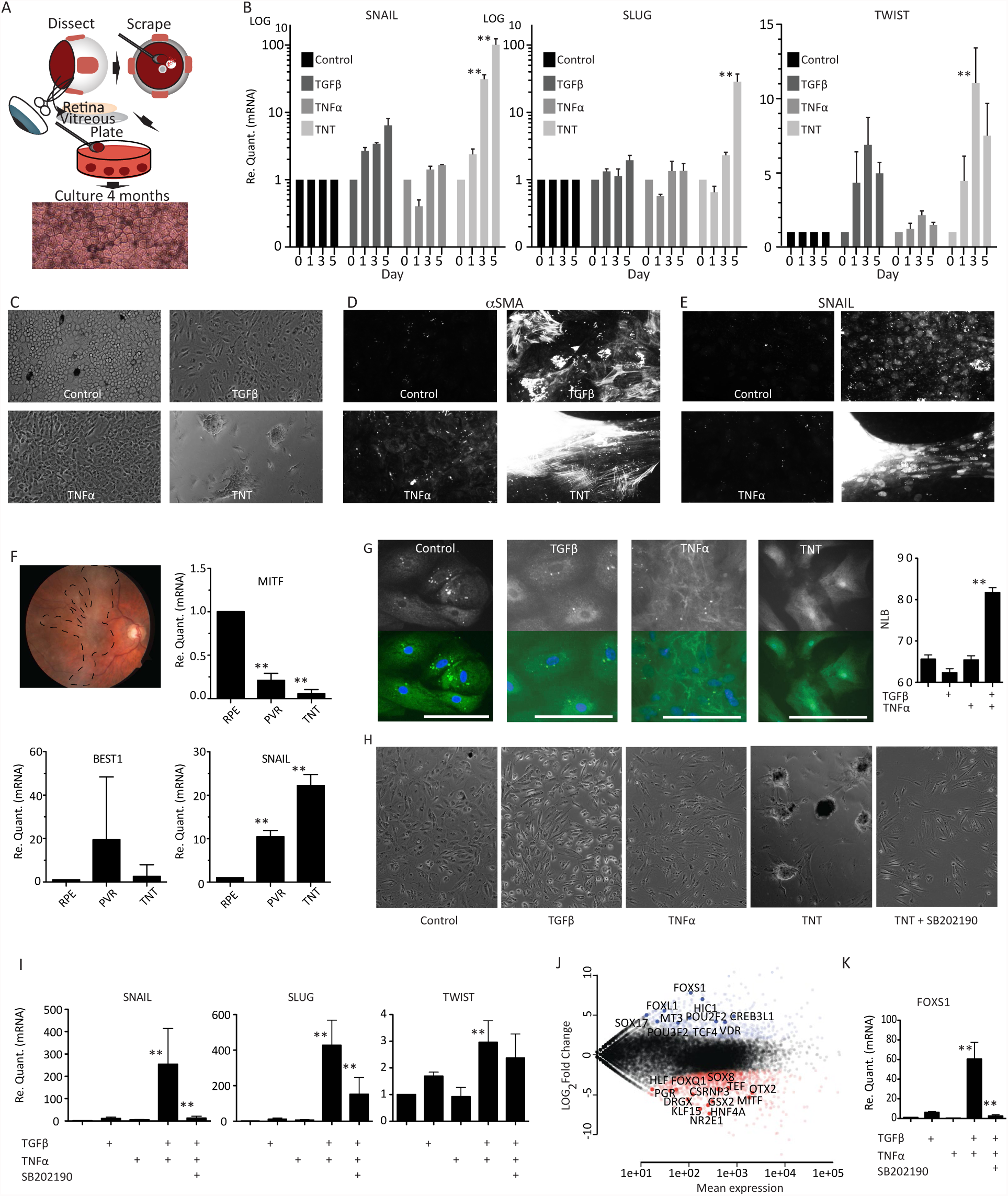
*TGFβ1+TNFα* co-treatment of cobblestone RPE induce s PVR-like EMT 3D masses. A) Schematic of the method to obtain pure cultures of RPE. B) Time course of *SNAIL*, *SLUG* and *TWIST* expression in control (cobblestone RPE, vehicle treated) and when treated with 10ng/ml of *TGFβ1* and/or *TNFα*. (C-E) Comparison of RPE cells in control conditions or 5 days after treatment with 10ng/ml of *TGFβ1* and/or *TNFα*. C) Phase images D) Immunostaining with anti-*SNAIL* antibody. E) Immunostaining with anti-αSMA antibody. F) Fundus images of patient with PVR membranes (indicated by dotted line) and gene expression comparison between TNT RPE and PVR samples from two patients. G) Immunofluorescence of p38 localization Inset: DAPI:blue, p38: green (NL=nuclear luminescence. H) Phase contrast images of RPE at day 5 of treatment indicated. I) qPCR of *SNAIL*, *SLUG* and *TWIST* from RPE after 5 days of treatment indicated. J) Scatterplot of RNA-seq data from cobblestone RPE versus TNT RPE with transcription factors highlighted; blue= increased and red= decreased expression. K) qPCR of *FOXS1* transcript after treatment indicated. Graph bars indicate SEM. Image scale Bars = 50μm. ** = P-value ≤ 0.01.

In addition to EMT transcription factor expression, TNT treatment induced a profound change in cell morphology, produced three-dimensional (3D) cell masses (Fig. 1C), and activated expression of *αSMA* and *SNAIL* (Fig. 1D,E), markers reported in PVR membranes surgically dissected from patients ^7,8^. To corroborate these findings with PVR disease, we next compared 5-day TNT treated versus untreated (vehicle control) RPE cells from 4 different donors to PVR membranes removed from four different patients (Fig. 1F). The PVR membranes expressed MITF at significantly higher levels 0.21 ± 0.79, *P* < 0.01 (mean +/- S. E. M; *n=4*) compared to control RPE (normalized to one for each sample), and compared to RPE treated with TNT 0.055 ± 0.023, *P* <0.01 (*n=4*). *BEST1* expression in PVR membranes varied widely 19.48 ± 14.47, *P* >0.05, (*n=4*), not significantly different from control RPE or TNT treated RPE 2.63 ± 2.63, *P* > 0.05, (*n=4*). SNAIL expression was significantly increased in PVR membranes 12.5 ± 2.13, *P* < 0.01, (*n=4*), compared to control RPE normalized to one, consistent with SNAIL expression of RPE treated with TNT 22.51 ± 1.46, *P* < 0.01, (*n=4)*. Taken together, the *in vitro* TNT model induces changes in RPE similar to those observed in PVR membranes. We therefore used this model to examine the cellular changes TNT evokes in RPE.

One of the known signaling pathways downstream of both *TGFβ* and *TNFα*, and therefore a candidate for their synergistic effect in RPE is the *p38 MAPK* pathway. To determine whether this was the case for human RPE, we examined whether TNT treatment induces p38 nuclear localization in RPE cells, testing this on five different donor lines. Indeed, *p38* was found to be preferentially localized in RPE nuclei in the TNT conditions 81.69 ± 1.17, *P* <0.01 (n=*167 cells)*), compared to control 65.61 ± 1.02 (n=*144)*), *TGFβ* 62.32 ± .99, (n=*161)*) or *TNFα* alone 65.4 ± 1.03 (n= *125)*) (Fig. 1G). Inhibiting p38 with the small molecule SB202190 (10ng/ml) prevented 3D mass formation (Fig. 1H) and suppressed *SNAIL* 0.04 ± 0.0067, *P* < 0.01 (*n=7*) and *SLUG* 0.34 ± 0.09, *P* < 0.01, (*n=7*), but not *TWIST* gene expression compared to the TNT condition (Fig. 1I).

In order to evaluate which additional transcription factors (TFs) may be involved in RPE-EMT, we compared cobblestone RPE and RPE with TGFβ1 alone, TNFα alone or TNT for five days via RNA-sequencing using 2 different donor RPE lines. 3136 genes were found to change significantly between the control and TNT conditions (FDR=0.01), and of those, 95 were known transcription factors. *FOXS1* was the most changed TF between these conditions (Fig. 1J).

We then asked whether p38 activation, induced by TNT was important in inducing *FOXS1* expression in seven experiments using four different donor lines. When SB202190 (10ng/ml) was added with TNT treatment, *FOXS1* expression was suppressed from 60.5 ± 17.19 to 2.57 ± 1.29, *P* < 0.01, (mean+/- S.E.M., *n=7*) (Fig. 1K), implying *FOXS1* is downstream of p38. *FOXS1* is expressed in peri-endothelial cells of the testis vasculature ^9^, in the aorta ^10^ and during the migration of neural crest derived sensory neurons ^11^, however it has not previously been linked to EMT. Hence, we have identified key steps along a cascade of changes during RPE transformation from an epithelial to mesenchymal phenotype. To further elucidate the events occurring in RPE EMT, we examined accompanying changes in the epigenome.

Changes in gene expression patterns during cell fate transitions are associated with changes in the epigenomic status at cis-regulatory elements. In particular, many distal enhancers are activated or decommissioned in a highly cell state-specific manner and this is accompanied by, respectively, gain or loss of certain chromatin marks (reviewed in ^12^). To systematically characterize epigenomic changes in RPE after EMT, we performed chromatin immunoprecipitation and sequencing (ChIP-seq) analysis with antibodies recognizing histone modifications marking active regulatory elements (*H3K4me1*, *H3K4me3*, and *H3K27ac*) using either control cobblestone RPE cells or those treated for 5 days with TNT. First, we examined genome-wide enrichments of *H3K27ac*, a histone modification associated with active enhancer and promoter states ^13,14^. Comparisons between untreated or TNT treated RPE revealed a major reorganization of *H3K27ac* patterns upon EMT, with a large number of sites gaining, and a smaller subset of sites losing *H3K27ac* in TNT conditions (Fig. 2A). Based on the relative enrichments between the two cellular states, we defined top *H3K27ac* sites preferentially enriched in TNT conditions (“TNT-specific”, highlighted in red) or untreated RPE (“RPE-specific”, highlighted in blue), and a similar number of sites at which *H3K27ac* enrichment changed least during treatment (“shared”, shown in orange) (Fig. 2A).

**Figure 2.**
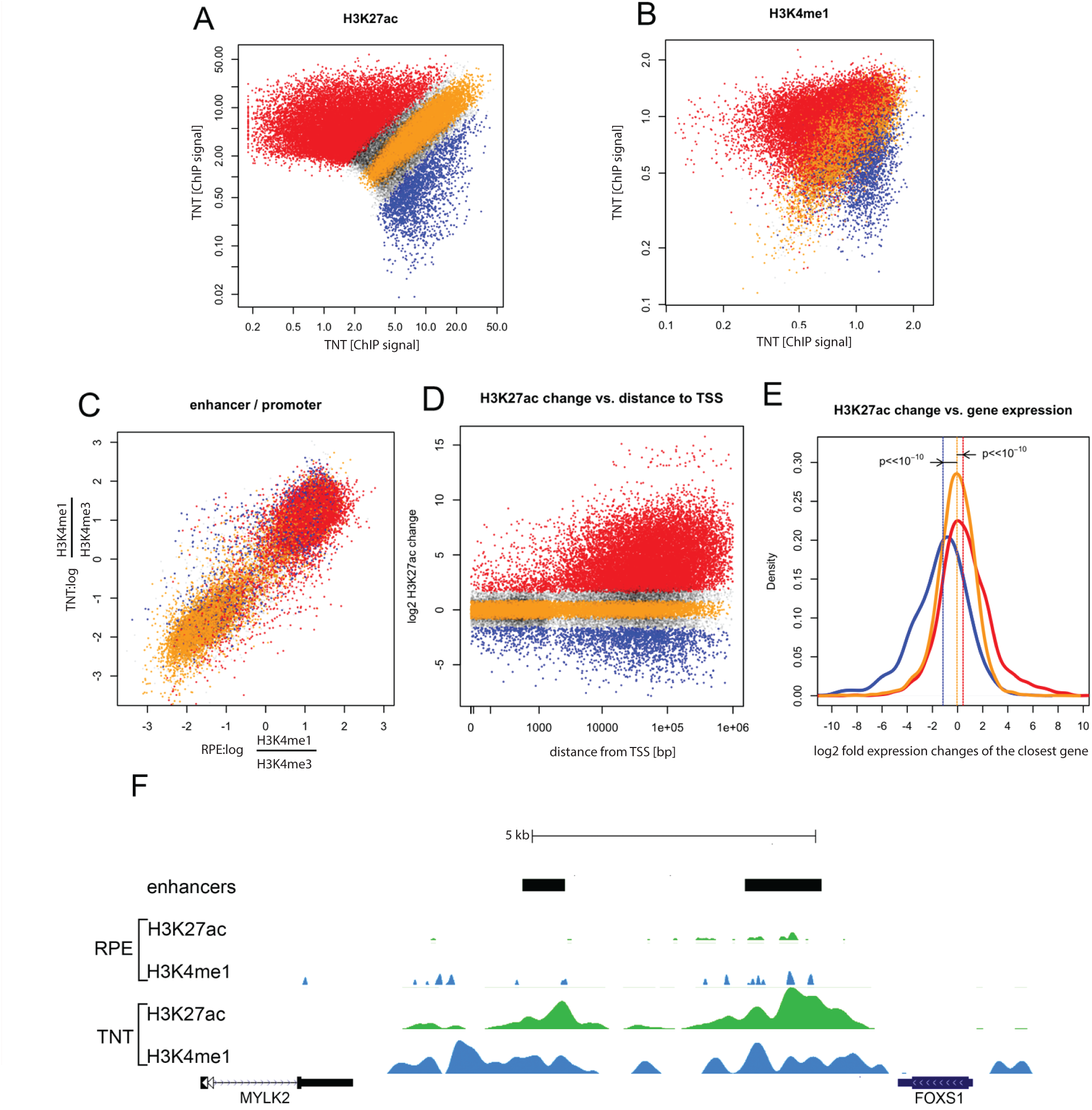
*TGFβ1*+*TNFα* co-treatment of cobblestone RPE induces massive epigenetic changes at enhancer elements. A) *TGFβ1* +*TNFα* co-treatment induced changes in *H3K27* acetylation: ordinate - normalized read density of *H3K27ac* ChIP-seq at chromatin features (putative enhancers and promoters) in cobblestone RPE cells, abscissa - normalized read density of *H3K27ac* ChIP-seq in *TGFβ1* +*TNFα* treated cells. Regions indicated in red have significantly upregulated ChIP signal upon *TGFβ1*+*TNFα* treatment (FDR<0.01), in blue downregulated (FDR<0.01), in orange with no change (FDR<0.1). B) Changes in the H3K4me1 ChIP signal are correlated with *H3K27ac* changes, regions are color coded according to the *H3K27ac* classes defined in the previous panel C) *TGFβ1*+*TNFα* co-treatment results in relatively few changes in promoter signatures. Plotted is the log ratio of *H3K4me1* versus *H3K4me3* ChIP density; RPE on the ordinate, treated cells on the abscissa. Negative values are indicative of a promoter-like chromatin signature at interrogated sites. Color-coding as in previous panels. D) Most changes in *H3K27* acetylation occur at sites distal from annotated transcription start sites. Plotted is the absolute distance to the closest annotated TSS versus the log2 fold change in the *H3K27ac* ChIP signal. E) Changes in *H3K27ac* are correlated with changes in gene expression. Plotted are the distribution of log2 fold expression changes for genes associated with distal elements that have upregulated *H3K27ac* (red), downregulated *H3K27ac* (blue) or were unchanged (orange). The differences are significant (Mann-Whitney-Wilcoxon test). F) Visualization of the histone modification changes at the *FOXS1* locus after *TGFβ1* +*TNFα* treatment in the UCSC browser; *H3K27ac* density in green, *H3K4me1* in blue. Location of two major FOXS1 enhancers is indicated by black bars.

We then analyzed *H3K4me1*, a histone mark associated with enhancer elements at the three classes of *H3K27ac* sites (as defined in Fig. 2D) and observed that changes in *H3K4me1* generally followed those in *H3K27ac*: TNT-specific *H3K27ac* sites had high levels of *H3K4me1* in TNT, but not in control RPE and *vice versa*, whereas shared *H3K27ac* sites were similarly enriched in both states (Fig. 2B). These observations suggested that major reorganization of *H3K27ac* patterns might occur at distal enhancer regions rather than proximal promoters. To confirm this, we examined changes in *H3K27ac* in relation to the *H3K4me1*/*H3K4me3* signal ratio, because relative enrichment of *H3K4me1* to *H3K4me3* can reliably distinguish putative distal enhancers from promoters (with enhancers having high *H3K4me1* and promoters *H3K4me3* enrichment) ^15^. We observed that sites that either gained or lost *H3K27ac* in TNT RPE (highlighted in red or blue, as defined in Fig. 2A) were generally characterized by the high *H3K4me1*/*H3K4me3* ratio, consistent with enhancer identity, whereas unchanged sites (orange) often had low *H3K4me1*/*H3K4me3* ratio, consistent with a large subset of the latter sites having promoter identity (Fig. 2C). In further agreement, dynamically changing *H3K27ac* sites were preferentially located more than 10 kb from the nearest transcriptional start site (TSS), whereas unchanged sites showed no such preference (and in fact were more enriched in TSS proximity) (Fig. 2D). Functional annotation of these dynamically changing putative enhancer regions revealed association of RPE-specific enhancers with retinal epithelium, visual perception and retinal degeneration categories, whereas enhancers active preferentially in TNT treated cells were enriched for annotations linked to vascular abnormalities (which with the EMT results, may suggest a role in vascular cell migration), cell growth and proliferation (Fig. 2. figure supplements 1-3). Taken together, our data show that EMT in RPE cells is associated with a global reorganization of chromatin landscapes that occurs preferentially at distal enhancer elements.

Next, we associated *H3K27ac* regions within each class defined in Fig. 2A with nearby genes and investigated the relationship with changes in expression observed during the RPE to TNT transition (Fig. 2E). We observed that genes associated with RPE-specific *H3K27ac* were commonly downregulated during the transition, whereas genes associated with TNT-specific *H3K27ac* were commonly upregulated in TNT treated cells as compared to cobblestone RPE (Fig. 2E). Loci encoding TFs that changed expression during the transition (denoted in Fig. 1J), also typically underwent reorganization of enhancer *H3K27ac* patterns, as exemplified by the *FOXS1* locus, which gains high levels of *H3K27ac* and *H3K4me1* at nearby putative enhancer regions (Fig. 2F).

These transcriptomic and epigenomic analyses predict that *FOXS1* may be a key TF involved in orchestrating RPE EMT. Consistent with this hypothesis, *FOXS1* was expressed much more in the PVR samples, 49.66 ± 20.28 fold, *P* < 0.01, (mean+/- S.E.M., *n=4* donors) compared to cobblestone RPE (Fig. 3A). We then asked whether *FOXS1* is upstream or downstream of the canonical master EMT regulators *SNAIL*, *SLUG* and *TWIST*. *FOXS1* shRNA inhibited the increase in *SNAIL* 0.07 ± 0.026, *P* < 0.01, (*n=4*) and *SLUG* 0.24 ± 0.12, *P* < 0.01 (*n=4*) during TNT treatment (Fig. 3B) and prevented 3D mass formation (Fig. 3C). Overexpression of *FOXS1* for 5 days induced expression of *SNAIL* 23.88 ± 4.2, *P* < 0.01 (Mean+/-S.E.M., *n=3* donors), *SLUG* 1.94 ± 0.17, *P* < 0.01 (*n=3*) and *TWIST* 2.98 ± 0.96, *P* < 0.01 (*n=3*) over RPE control (Fig. 3D), reduced expression of the RPE markers *MITF* 0.05 ± 0.01, *P* < 0.01 (*n=3*) and *OTX2* 0.26 ± 0.07, *P* < 0.01 (*n=3*), but increased *BEST1* expression to 13.79 ± 4.12, *P* < 0.01 (*n=3*) (Fig. 3E), changes consistent with those seen upon TNT treatment and in PVR patient samples. However, *FOXS1* overexpression alone did not induce 3D mass formation (Fig. 3F). *FOXS1* is therefore necessary and sufficient to change *SNAIL* and *SLUG* gene expression.

**Figure 3.**
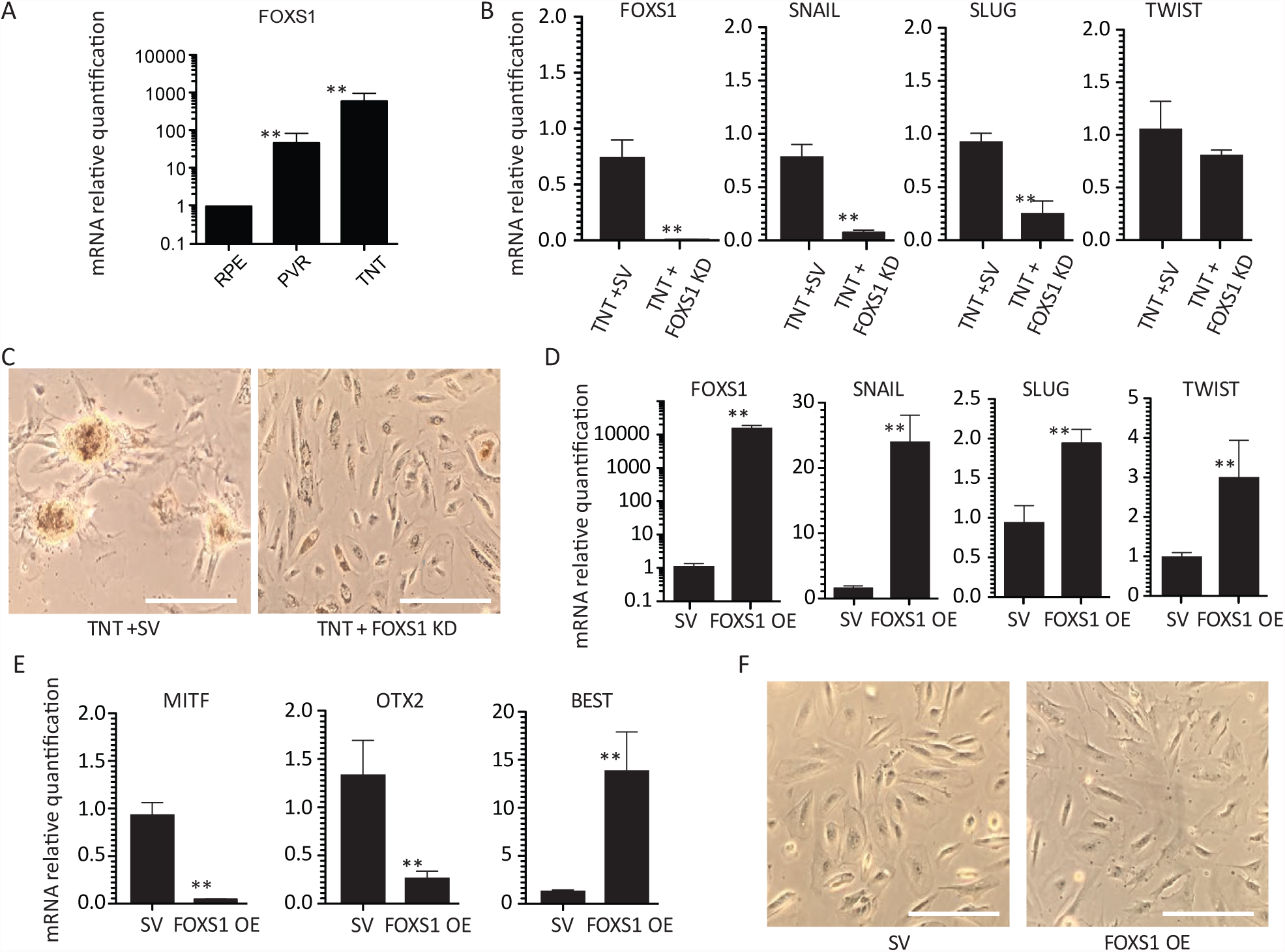
*FOXS1* is necessary and sufficient to drive *SNAIL* and *SLUG* expression in RPE. A) qPCR gene expression comparison between TNT-RPE and PVR samples from two patients. B) qPCR of RPE after 5 days of TNT and either a scrambled, or *FOXS1* shRNA lentivirus treatment and C) corresponding color images. qPCR gene expression comparison of RPE with either a scrambled or FOXS1 overexpression lentivirus of EMT (D) and RPE (E) genes. F) Color images of RPE at day 5 after treatment with either a scrambled or *FOXS1* overexpression lentivirus. Image Scale bars=100mm. ** indicates P-value ≤0.01. Graph bars = SEM. SV=scrambled control vector.

The observation that *FOXS1* regulates canonical EMT transcription factor expression led us to ask whether it plays a role in EMT in other epithelia, including human mammary epithelium (HME) and human hepatic epithelium (HHE). HME cells treated with TNT showed increased expression of both *FOXS1* and *SNAIL* (Fig. 4A), but not *SLUG* or *TWIST* (data not shown). *p38* inhibition blocked the expression of both *FOXS1* from 8.44 ± 0.61 to 0.23 ± 0.1, *P* < 0.01 (mean+/-S.E.M., *n=3* donors) and *SNAIL* from 16.23 ± 8.59 to 0.91 ±0.29, *P* < 0.01 (*n=5*) in HME under TNT conditions (Fig. 4A). Knocking down *FOXS1* during TNT treatment in HME cells decreased *SNAIL* expression to 0.096 ± 0.16, *P* < 0.01 (*n=3*) compared to RPE control (Fig. 4A), while overexpression of *FOXS1* induced *SNAIL* expression by 3.35 ± 0.58 fold, *P* < 0.01 (*n=3*), which is also seen at the protein level (Fig. 4B). Therefore, TGFβ1 and TNFα signaling synergizes through *p38*, leading to an increase in *FOXS1,* which then induces an increase in *SNAIL* expression in HME. Similar studies in HHE cells (Fig. 4C) indicate that *FOXS1* is sufficient but not necessary to induce expression of *SLUG,* and necessary and sufficient to change *TWIST* expression.

**Figure 4.**
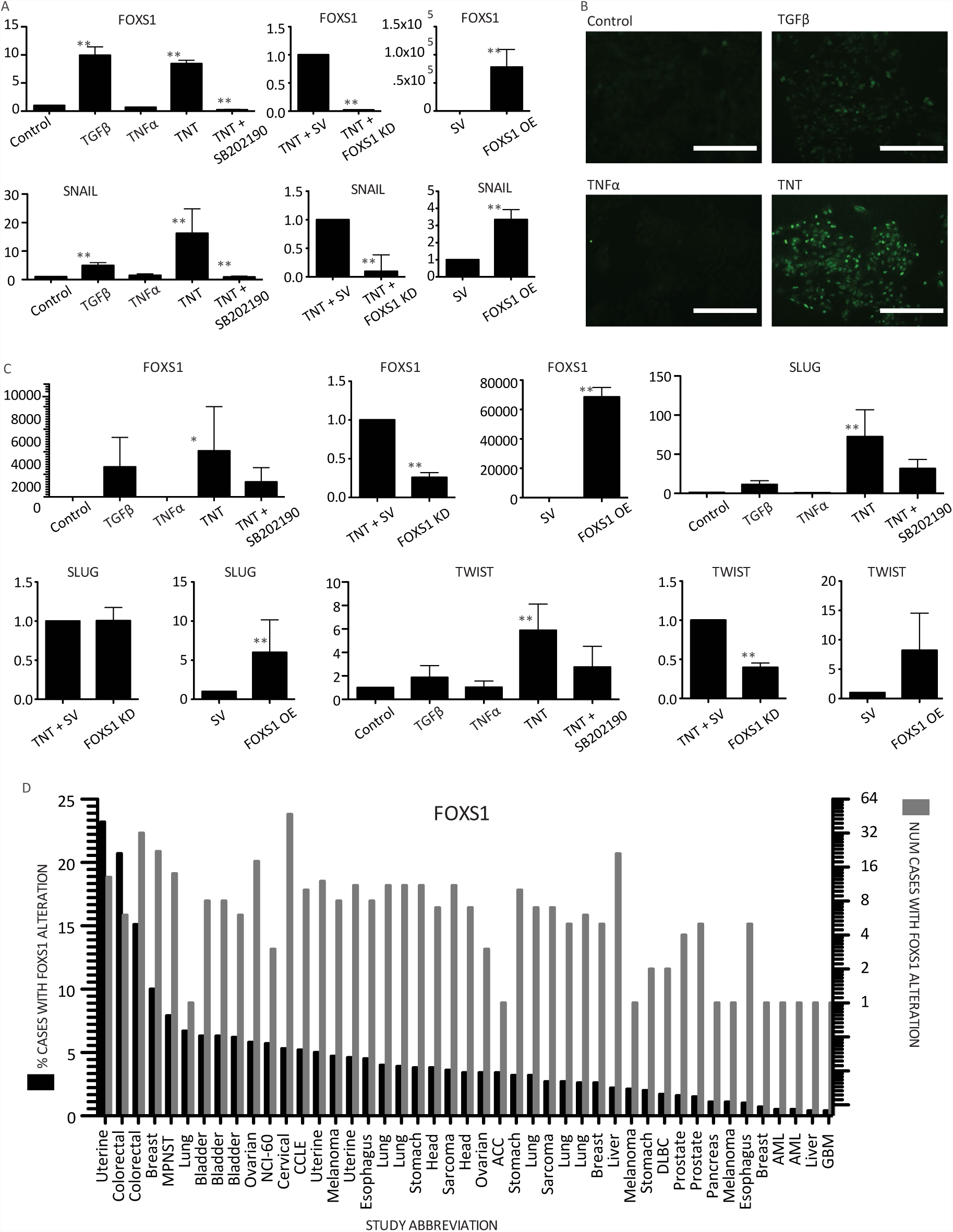
*FOXS1* in EMT in other epithelia and cancer. A) qPCR of HME cells treated with *TGFβ1* and/or *TNFα* with and without p38 inhibitor SB202190; Lentivirus treatment including a scrambled, or *FOXS1* shRNA in HME cells with TNT; lentivirus treatment of HME cells with either a scrambled or *FOXS1* overexpression construct. B) Immunofluorescence of HME cells treated with *TGFβ1* and *TNFα* stained with anti-SNAIL antibody. C) qPCR of HHE cells treated with *TGFβ1* and/or *TNFα* with and without p38 inhibitor SB202190; Lentivirus treatment including a scrambled, or *FOXS1* shRNA in HHE cells with TNT; lentivirus treatment of HHE cells with either a scrambled, or *FOXS1* overexpression construct. D) Cross cancer analysis shows *FOXS1* alterations are present in 47 of 92 studies examined (See Sup Table 1). Graph scale bars = SEM. Image scale Bars = 200μm. * = P-value ≤ 0.05. ** = P-value ≤ 0.01.

Since EMT is hypothesized to play a role in the metastasis in many cancers, we examined the incidence of *FOXS1* changes in several cancer studies ^16,17^. Out of 91 whole genome sequencing tumor studies analyzed, 47 included at least one case of multiple copies of the *FOXS1* gene (Fig. 4D), 14 had greater than 5% of cases and in one uterine cancer study, 23% of cases had multiple copies of *FOXS1*. Based on our observations of a central role in EMT, we hypothesize that *FOXS1* may be involved in the metastatic invasion of cancer cells.

## Discussion

EMT is a complex, multifactorial process resulting in profound changes in cell phenotype and behavior. The molecular mechanism underlying EMT is still not completely resolved. In this study we defined the global transcriptomic and epigenomic changes associated with EMT in human RPE cells. Subsequent analysis elucidated factors associated with EMT-linked pathology in several cell types, and identified FOXS1 as a key driver of the process, and a novel upstream factor which regulates known master EMT transcription factors.

Here we show that *TGFβ1* or *TNFα* treatment alone cause slight alterations in RPE morphology, but together they cause a dramatic induction of EMT, during which the RPE cells lose their normal cobblestone configuration and grow into 3D masses that lose key RPE features and gain mesenchymal characteristics. The synergistic action of these factors in stimulating EMT has been reported for non-neural epithelia, including from the lung ^18^ and colon ^19^, and in the latter case, they do so via *p38*-*MAPK* signaling, implying a common mechanism is shared by neural epithelia. TNT treatment resulted in numerous changes in the RPE epigenome, with a bias towards gene activation. The changes in active chromatin signatures occurred largely at distal enhancers, while promoters were much less affected, consistent with enhancers serving as key mediators of changes in cell states and being utilized in a highly dynamic manner (reviewed in 12). GO analysis of the significantly changing enhancers highlighted those associated with extracellular remodeling, vasculature, wound healing, and cell proliferation, which align with the stimulation of cell proliferation and migration apparent in the cultures. The changes occurred within 5 days of treatment, demonstrating that stable cobblestone RPE can rapidly disassemble and form 3D masses in the presence of stimulating environmental factors.

Single nucleotide polymorphisms which confer a significantly increased risk of PVR have been located at both TGFβ1^5^ and TNFα^4^ genetic loci. Moreover, TGFβ1 and TNFα have both been reported at increased levels in the vitreous of patients with epiretinal membranes ^3,20^ and TGFβ1 in particular is associated with their maturation into contractile masses that lead to retinal detachment and severe vision loss ^21^, providing the impetus for use of Rho-kinase inhibitors to prevent the myosin-based contractions. While inhibition of contraction of already formed epiretinal membranes is a valuable therapeutic approach, by determining the causes of RPE EMT, our goal is to identify targets that inhibit the production of these membranes at an earlier stage of the disease.

Within 5 days of EMT induction, RPE cells increase expression of 95 transcription factors that may play roles in activating the pleiotropic phenotypic changes. Of these, the most dramatic increase occurs in *FOXS1*, which is not detectable in the control cobblestone RPE. Changes in histone marks surrounding the *FOXS1* gene locus were consistent with its activation. We have corroborated expression of *FOXS1* in patient PVR samples. Most importantly, knockdown of *FOXS1* inhibited production of 3D masses, and was found to be both necessary and sufficient to drive the canonical EMT master regulators SNAIL and SLUG in human RPE cells. These observations implicate *FOXS1* induction as an early, causal factor in pathological RPE EMT (modeled in Figure 5), and raise questions about the specific targets of FOXS1, which will be the focus of future studies. Our finding that p38 inhibition can prevent the increase in FOXS1 raises the possibility of an early intervention in this pathological process.

**Figure 5.**
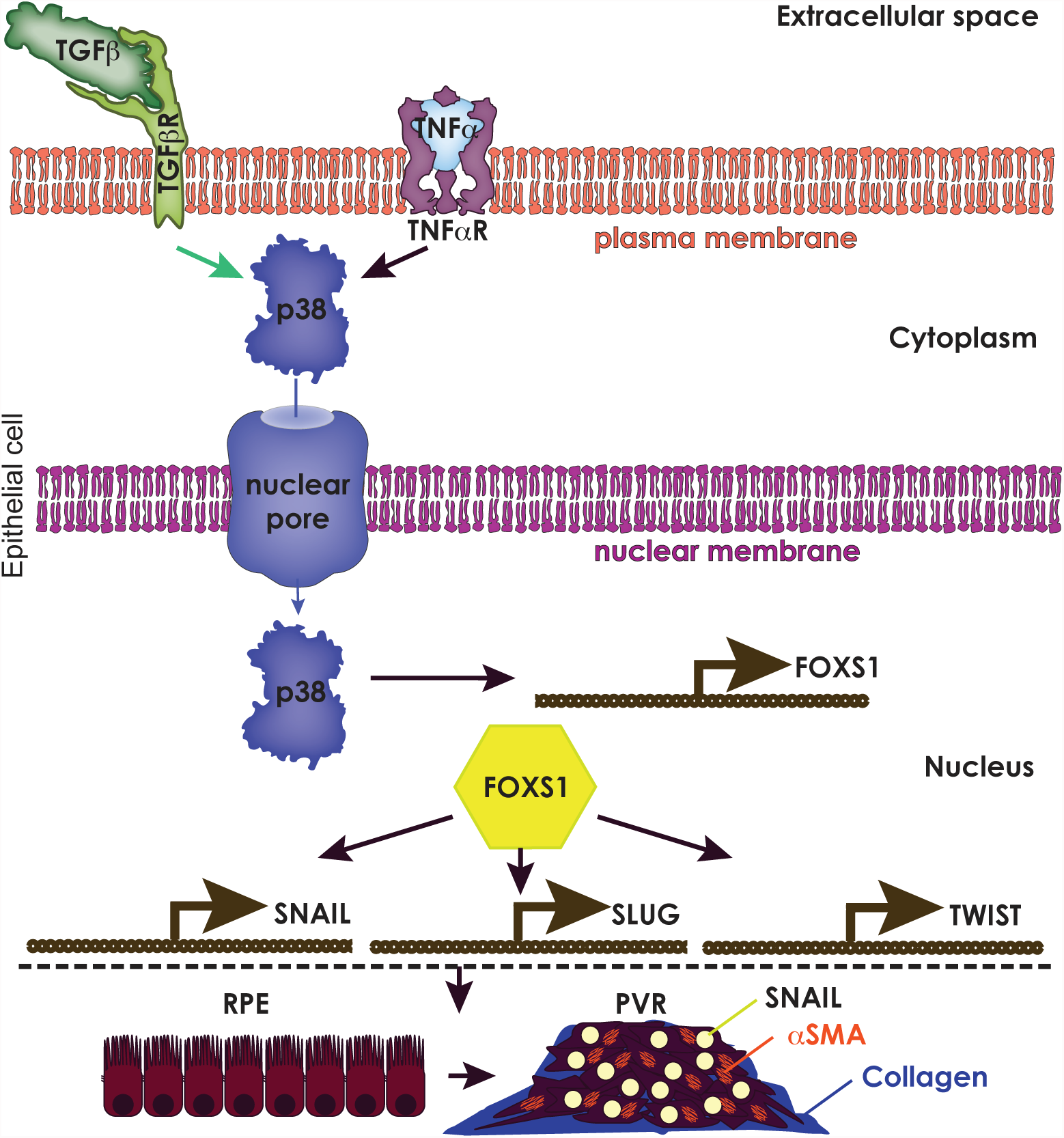
Model of the pathway of synergistic action of *TGFβ* and *TNFα* in transforming RPE into 3D contractile tissue. *TGFβ* and *TNFα* bind to their respective receptors leading to a co-activation of p38, which leads to p38 translocation into the nucleus and transcriptional activation of *FOXS1*. *FOXS1* expression leads to *SNAIL*, *SLUG*, and *TWIST* expression, driving an epithelial to mesenchymal transition in RPE.

In addition to its action in RPE cells, we demonstrated that *FOXS1* can act as a master transcription factor in EMT in other cells, as it is upstream of the canonical EMT transcription factors *SNAIL*, *SLUG* and *TWIST* in two non-neural epithelial cell types tested. The *FOXS1* gene can be hypomethylated in adrenocortical neoplasms ^22^ and demonstrates increased copy number across a variety of cancers. Given that *FOXS1* is not expressed in normal RPE nor in other adult epithelia, its expression and involvement in pathological EMT indicate it may be a valuable therapeutic target across several tissue types.

## Acknowledgements

We thank Dr. Barbara Corneo for manuscript review, Carol Charniga and Dr. Marie Fernandes for their invaluable help with eye dissection and cell culture, and the eye donors and their families for the generous donation of retinal tissue. This study was supported by a National Eye Institute (NEI; Bethesda, MD, USA) extramural grant (1R01EY022079; ST) and by NYSTEM through a retinal stem cell consortium grant (N11C012) and by the Icahn School of Medicine at Mount Sinai (TAB) and by a gift from Dr. Heinrich Medicus to the Regenerative Research Foundation (S.T. and J.H.S).

## Author contributions

TAB, ST, JHS and JW conceived the experiments, and helped with data analysis, interpretation and manuscript writing. TAB performed the experiments, analyzed data, made figures, and wrote the manuscript. TAB and QW produced shRNA and overexpression FOXS1 lentiviruses, RS prepared RNA and DNA for sequencing, TS and NB analyzed sequencing data and made figures. ARI performed 1 ChIP-seq and data analysis. TAB wrote the initial manuscript draft, which was edited primarily by ST and JW and additionally by TS, NB, RS, QW, JHS.

## Author Information

Data deposition with URL and accession database numbers

## Methods

### Human adult RPESC culture

Human globes from donors aged between 51 and 89 yrs were obtained from the National Disease Research Interchange, Philadelphia, PA., the Eye-Bank for Sight Restoration, Inc., New York, NY, and the Lions Eye Bank, Albany, NY. A detailed eye dissection protocol was previously published ^23^. Globes were obtained within 40 hours of death, RPE sheets were isolated and plated on tissue culture plates coated with 10μg/ml placental ECM (Corning) or Synthemax II (Corning) in RPE medium ^23^, supplemented with 10% FBS and 10mM Nicotinamide, which was changed 3 times a week. After the first week, FBS was reduced to 2%.

### EMT model

Passage 0 adult human RPE cultures (ahRPE) were trypsinized using 0.25% Trypsin for 15 minutes, washed and replated at 3 x10^4^ cells per 1.9mm^2^ per well of a 24-well plate in DMEM/F12 with 5%FBS, L-Glutamine, Na-Pyruvate, NEAA, Pen/Strep. After 24 hours, 10ng/ml TGFβ1 or TNFα, or both (TNT) were added to induce EMT versus vehicle as control, and the cultures were maintained in this medium (with feeding every other day) for 5 days.

### Immunohistochemistry

ahRPE on 24 well-size transwell inserts (Corning) were fixed with 4% paraformaldehyde for 10 minutes, rinsed 3 times with phosphate buffered saline (PBS), permeabilized with 0.01% saponin and blocked with normal goat serum (5%) in 1% BSA in PBS for 1 hour. Primary antibody (Table 1) was added overnight at 4^o^C, then incubated with the corresponding Alexa Fluor conjugated secondary antibodies (1:1000) (Life Technologies Alexa Fluor 647 goat anti-mouse IgG (H+L) (Cat#: A-21237), Alexa Fluor 647 goat anti-rabbit IgG (H+L) (Cat#: A-21244), at room temperature for 45 minutes. Cells on transwell inserts were then mounted on glass slides with Prolong gold (Life Technologies) and imaged by confocal microscopy (Leica). DAPI fluorescence was used to demarcate the nucleus in experiments measuring nuclear p38. P38 fluorescence intensity was then measured exclusively in the DAPI demarcated region based on a 0-256 scale using ImageJ analysis software. The fluorescence intensity of all nuclei was averaged across experiments and compared between conditions using a student’s t-test.

### qPCR

RNA was extracted (Qiagen) and reverse-transcribed (Superscript III Reverse Transcriptase kit) then qPCR was performed according to manufacturer instructions (Power SybrGreen PCR Master Mix) using primers in Table 2. Data was recorded as the cycle by which sybr fluorescence detection threshold is reached during the exponential increase phase. All cycles for queried genes were normalized to the housekeeping gene cyclophilin to control for cDNA quantity variation.

**Table 2.**
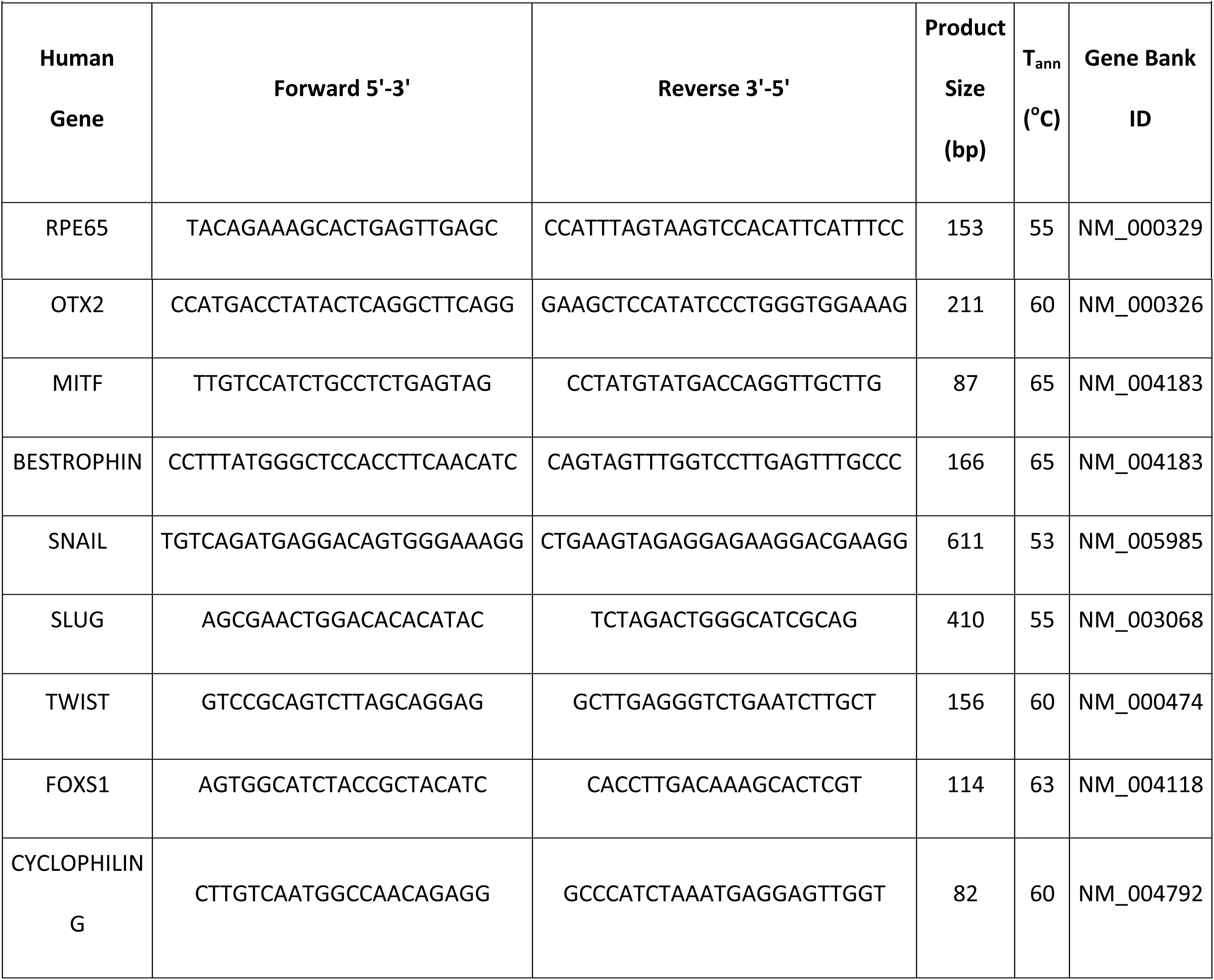
List of primers used for Real Time PCR on adult human RPE

### Lentiviral vector production and viral packaging

shRNA plasmids: the hairpin oligonucleotides (Scrambled control: TTCTCCGAACGTGTCACGT; FOXS1 3’ UTR: GCCAATAAAGCCATGTGAT) were inserted into the FUGW-H1 lentiviral construct as previously described ^24,25^. To package the lentivirus, the constructs were co-transfected with pCMV-VSVG and pCMV-dvpr into 293FT cells. Supernatant was harvested 2 and 3 days later and concentrated by ultra-centrifugation. For overexpression constructs, the open reading frame (ORF) of each gene was PCR-amplified, sequence-verified, and cloned into a modified version of FUGW. An IRES-eGFP was included for visualization. Lentiviruses were used at 10 MOI for cell transduction.

### ChIP-seq

ChIP assays were performed from 10^7^ RPE cells per experiment, according to a previously described protocol with slight modification ^26^. Briefly, RPE cells at passage 0 cultures were treated with TGFβ1, TNFα or TNT conditions for 5 days. The cells were then with 1% formaldehyde for 10 minutes at room temperature to crosslink, and the reaction was then quenched by adding glycine at a final concentration of 0.125 M. Cells were dissolved in lysis buffer and chromatin was sonicated to an average size of 0.5–2 kb, using Bioruptor (Diagenode). 5–7.5 μg of antibody was added to the sonicated chromatin and incubated overnight at 4°C. Subsequently, 50μl of protein G Dynal magnetic beads were added to the ChIP reactions and incubated for 4–6 hr at 4°C. Magnetic beads were washed and chromatin eluted, followed by reversal of crosslinks and DNA purification. ChIP-seq and input libraries were prepared according to Illumina protocols and sequenced using the Illumina HiSeq. To identify the approximate positions of regulatory elements from histone modification ChIP-seq profiles alone, we calculated kernel density estimate tracks with bi-modal kernels ^27^ and identified peaks in the product of the combined H3K27 and H3K4 signals. To analyze changes in histone modifications, we calculated the read coverage for each sample over a combined set of detected peaks and performed a differential analysis with DESeq2.

### RNA-seq

RPE cells from two different donors were treated with TGFβ1, TNFα or TNT or vehicle control as described above for 5 days, then the RNA was extracted with Trizol (Invitrogen), following the manufacturer’s recommendations. 10ug of total RNA was subjected to two rounds of purification using Dynaloligo-dT beads (Invitrogen) then fragmented with 10× fragmentation buffer (Ambion) and used for first-strand cDNA synthesis, using random hexamer primers (Invitrogen) and SuperScript II enzyme (Invitrogen). Second strand cDNA was obtained by adding RNaseH (Invitrogen) and DNA Pol I (New England BioLabs). The resulting double-stranded cDNA was used for Illumina library preparation and sequenced with the Illumina HiSeq. Reads were mapped with tophat v2.0.14 to the gencode 23 transcriptome model and the read coverage was calculated with htseq. Differential analysis was performed with DESeq2.

